# Instant three color multi-plane fluorescence microscopy

**DOI:** 10.1101/2021.05.07.443091

**Authors:** Ingo Gregor, Eugenia Butkevich, Jörg Enderlein, Soheil Mojiri

**Affiliations:** III. Institute of Physics – Biophysics, Georg-August-University, 37077 Göttingen, Germany; Cluster of Excellence “Multiscale Bioimaging: from Molecular Machines to Networks of Excitable Cells” (MBExC), Georg-August-University, 37077 Göttingen, Germany

## Abstract

One of the most widely used microscopy techniques in biology and medicine is fluorescence microscopy, offering high specificity in labeling as well as maximum sensitivity. For live cell imaging, the ideal fluorescence microscope should offer high spatial resolution, fast image acquisition, three-dimensional sectioning, and multi-color detection. However, most existing fluorescence microscopes have to compromise between these different requirements. Here, we present a multi-plane multi-color wide-field microscope that uses a dedicated beam-splitter for recording volumetric data in eight focal planes and for three emission colors with frame rates of hundreds of volumes per second. We demonstrate the efficiency and performance of our system by three-dimensional imaging of multiply labeled fixed and living cells.

## 1. Introduction

The ultimate goal of fluorescence microscopy is to record three-dimensional images of cellular structures with maximum spatial and temporal resolution and high signal-to-noise ratio, ideally in a robust and easy to use manner. While super-resolution microscopy has pushed the limits of spatial resolution down to a few nanometers, [1–8] it could do so only by compromising on simplicity of the utilized hardware or data analysis approaches, on image acquisition speed, and/or on the ability of imaging several structures/molecules with different emission colors at the same time, i.e, multiplexing [9–11]. However, (three-dimensional) imaging speed and multiplexing are pivotal for following multiple cellular processes *in vivo* in three dimensions and with sufficient temporal resolution.

To achieve maximum acquisition speed of whole volume images, several multi-plane wide-field imaging modalities have been recently developed. In 2013, Abrahamsson and coworkers introduced the first multiplexed 3D wide-field epi-fluorescence microscope using a chromatically corrected grating that splits the emission light into several detection channels, each corresponding to a different focal plane and to two emission colors [12]. The authors of ref. [13] successfully used such a grating-based multi-plane microscope even for single-molecule localization microscopy, although the signal-to-noise ratio goes increasingly down with increasing number of simultaneously imaged planes. Later, the Abrahamsson group improved the chromatic aberration correction and optical efficiency of this microscope by incorporating two gratings and two aberration-compensating prisms, one grating/prism pair for one color [14]. Although the development of these systems is a formidable technological achievement, their technological complexity (complex micro-optical phase gratings and chromatic-aberration correcting multifaceted prisms) and demands on alignment has prevented its widespread application so far. Furthermore, up to now such a system was realized only for a maximum of two spectral channels.

As an alternative, Leutenegger and colleagues developed a glass-based beam splitter device that allows for multi-plane imaging similar to the diffractive grating approach by Abrahamsson, but in a much simpler manner and with tremendously reduced chromatic aberration. While the first setup of this beam splitter still used an array of individual 50/50 beam splitters and mirrors, it was later simplified into a single glass-piece which offers maximum simplicity in implementation, maximum mechanical stability, and nearly no chromatic aberration over the spectral range of 500 nm to 700 nm [15–18].

In the present paper, we expand this multi-plane beam-splitter system with an additional spectral beam-splitter to achieve simultaneous three-color imaging in eight focal planes with a volumetric image acquisition rate of up to several 100 volumes per second in three spectral windows. Besides providing high imaging speed and color multiplexing, our system is straightforward to implement and to use, both in hardware and in data analysis.

## 2. Methods

### 2.1. Setup

A schematic of our three-color multi-plane imaging setup is shown at the top of Fig. 1. Fluorescence excitation is done with three light sources, a 100mW laser at 473 nm (CNI), a 200mW laser at 561 nm (CNI), and a 140mW laser at 637 nm (Obis, Coherent). All three beams are aligned to a common path, spatially filtered, and expanded. The collimated beam is guided into the illumination path of a conventional inverted epi-fluorescence microscope (IX-71, Olympus) equipped with high-NA water immersion objective (UPLSAPO 60× 1.2 NA W, Olympus). A schematic of the illumination configuration is shown in Fig. S1 of the Supplementary Information (SI).

**Fig. 1.**
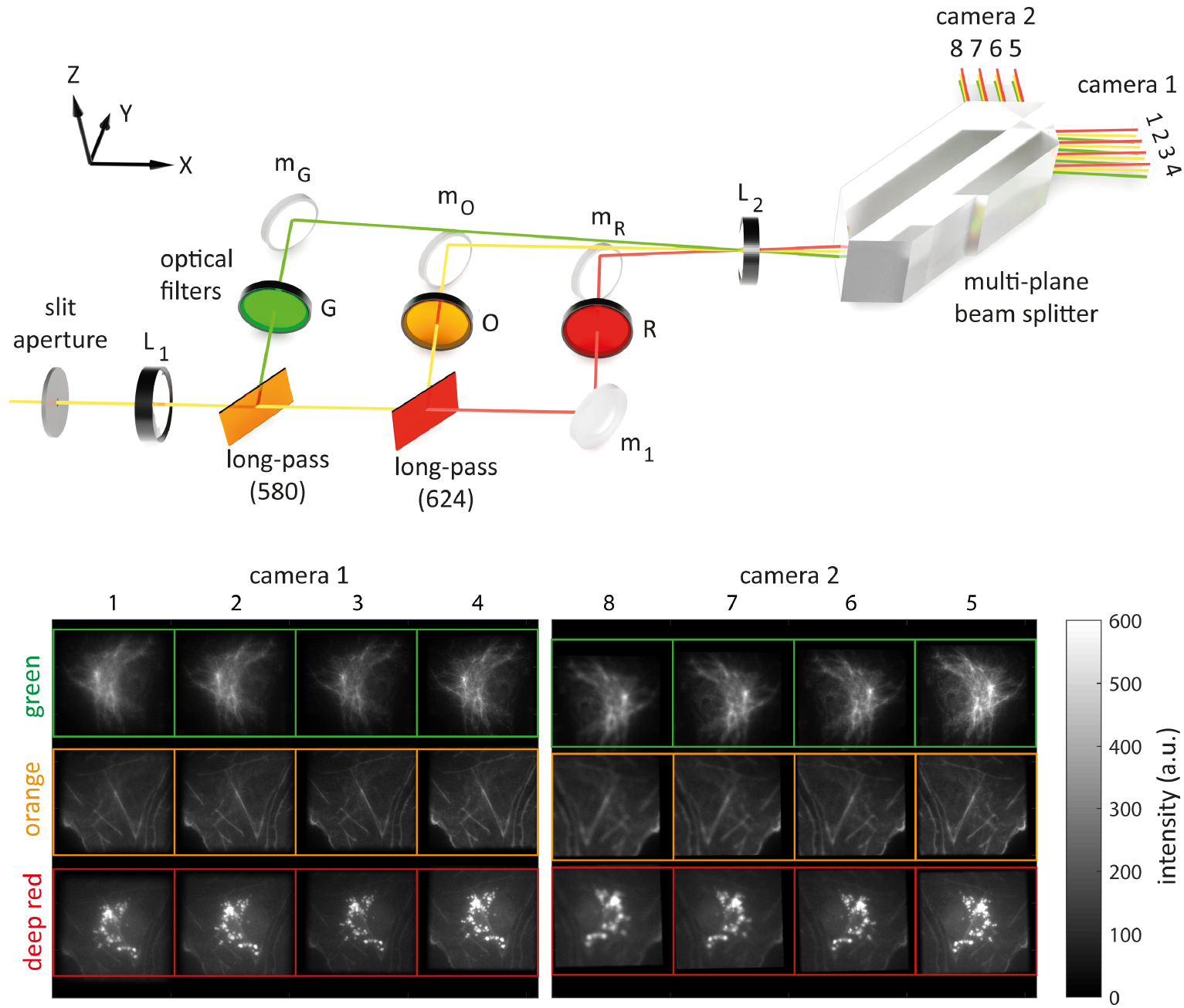
Top: Detection path of the 3-color multi-plane fluorescence microscope. An adjustable slit aperture crops the light at the microscope’s side-port, thus defining the field-of-view. Lenses L_1_ and L_2_ comprise a 4f optics with magnification 1.33 that relays the fluorescence light to the sCMOS cameras 1 and 2. Between the lenses, the light sequentially passes two dichroic mirrors. The resulting three color channels of green (*λ* < 580 nm), orange (580 < *λ* < 624 nm), and red (*λ* > 624 nm) are further filtered with emission bandpass filters G (513/17 nm), O (593/40 nm), and R (692/40 nm), respectively. An off-axis tilt of the dichroic mirrors deflects the propagation direction of the green and deep-red color away from the horizontal plane. Next, the multi-plane prism generates eight images with uniformly increasing optical path length, corresponding to eight distinct focal planes inside the sample. Bottom: Raw eight-plane three-color images of a fixed fluorescently labeled COS-7 cell. Different focal planes are shown along the horizontal direction, different colors along the vertical direction. The different colors correspond to labeled vimentin, actin, and mitochondria with the corresponding peak emission wavelengths at 520 nm, 593 nm, and 692 nm, respectively. The numbers above the images show the axial order of the image planes (1 is closest to sample surface, 8 is farthest).

The fluorescence detection path comprises a color-splitting unit, and a multi-plane beam-splitting unit. After passing the first lens L_1_, a custom-built three-color beam splitter splits the fluorescence light into three spectral channels along the vertical direction: a green channel (G, 450 nm to 580 nm), an orange channel (O, 580 nm to 624 nm), and a deep-red channel (R, 650 nm to 700 nm). To spatially separate these color channels on the camera chip, each color is reflected under a slightly different vertical angle: The middle color channel, i.e. the orange beam, propagates in the plane, while the green and deep-red beams are tilted by angles ±*θ*. The angle *θ* is determined by the required vertical offset Δ*y* of the images on the camera and the focal length *f*_2_ of the final tube-lens as *θ* ≥ arcsin (Δ*y*/*f*_2_). In our case we have Δ*y* = 3.5 mm and *f*_2_ = 200 mm, resulting in *θ* ≈ 1°. Note, that the length of the optical path is the same for all three color channels. The mirrors *m*_G_, *m*_O_, and *m*_R_ align the beams of the three channels to a common vertical plane. Here, the orange and green beams pass over the mirrors *m*_O_ and *m*_R_, respectively. A detailed schematic of the spectral beam-splitting unit with inter-component distances is provided in in Fig. S2 of the SI.

Next, all three colors are passed through a tube lens (L_2_) and a commercially available multi-plane beam splitter (patent EP3049859A1, Scoptonic imaging technologies, Torun, Poland). This multi-plane beam splitter is geometrically designed in such a way that it splits an input beam, through multiple reflections, into two sets of four neighboring output beams with increasing optical path length [15, 19]. The focal lengths of lens L_1_ and L_2_ are chosen in such a way that the final magnification increases by a factor 1.33. After the multi-beam splitter, the two sets of four images are recorded by two sCMOS cameras (ORCA-Flash 4.0V2, Hamamatsu), where the second camera is shifted, as compared to the first camera, by an additional distance 4*d*/*n* ~ 9.16 mm away form the prism (here, *n* = 1.46 is the refractive index of the prism material). In this way, all eight images recorded by the two cameras correspond to eight equidistantly placed focal planes in sample space, with an optical path difference of Δ*l* = *d/n* between them. An example of a recorded set of 8 × 3 images (8 focal plane times 3 colors) is shown in Fig. S3 of the SI. The field aperture prevents lateral overlap of neighboring images on the camera chips.

To determine the precise distances between focal planes, and their relative light-collection efficiencies, a stack of *z*-scan images over an axial range of 7 μm in steps of 100 nm of immobilized tetra-spectral fluorescent beads was recorded (TetraSpeck™ carboxylate micro-spheres, 0.1 μm, T7279, Thermo Fisher Scientific). Fig. S4 in the SI shows one such image of fluorescent beads. Fluorescence excitation was done using the 473 nm and 561 nm lasers. The average bead brightness in each plane over the *z*-scan can be well fitted with a Gaussian function, see Fig. 2. From these 24 fits, the axial shift of each plane and the relative collection efficiency of each plane can be determined. Notice that the latter is not identical for all focal planes, due to slight imperfections of the multi-plane beam-splitter prism.

**Fig. 2.**
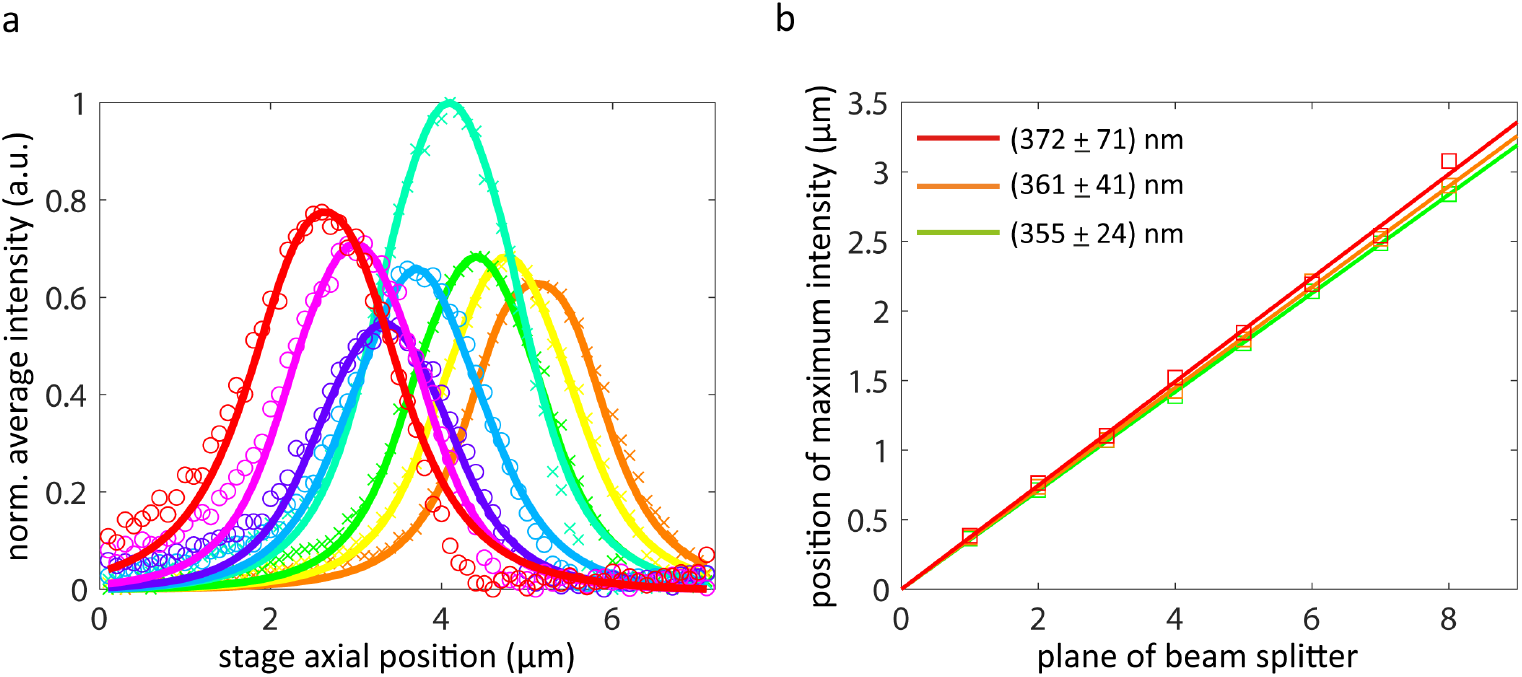
Brightness and inter-plane distance calibration. (a) Normalized average intensity of image planes in the green channel as a function of axial scan position (scan step equals to 100 nm). Solid lines represent Gaussian fits, crosses (camera 1) and open circles (camera 2) are measured data. (b) Linear fits of z-scan positions of intensity maxima as a function of image for each color channel. These fits give an inter-plane distance of 355 ± 24 nm for the green channel, of 361 ± 41) nm for the orange channel, and of 372 ± 71 nm for the deep red channel.

As can be seen from the right panel in Fig. 2, the maxima of the Gaussian fits for each image plane follow a clear linear dependence in plane number, from which the uniform inter-plane distance for each color channel can be determined. These distances are slightly different for the three color channels (less than 5%) due to chromatic effects. The color-dependent differences of inter-plane distance accumulate to maximally 140 nm over the full axial depth range of ≈ 2.5 μm over entire spectral range from 520 nm to 685 nm. This is consistent with analytical color aberration values (≈ 220 nm) over a range from 500 nm to 700 nm obtained by ray tracing analysis of the multi-plane beam-splitter prism (see SI in Ref. [15]).

The spectral separation of the three-color beam splitter is not perfect. The green emission is recorded with 14.68% in the orange channel and with 0.97% in the deep red channel. Similarly, orange emission is recorded with 13.38% in the deep red channel (see lower panel in Fig. 1). Finally, for determining the transmission efficiency of the three-color beam splitting system, we measured the intensity of 20 tetra-spectral beads in the orange channel with and without the three-color beam-splitter and long-pass filters in place. In this way, we found a transmission efficiency of ≈ 89%.

### 2.2. Image analysis

All image processing was done with custom-written Matlab routines. To generate a *z*-stack for each color channel, neighboring images (see colored rows in Fig. S4) are cropped, weighted by the corresponding collection efficiency obtained from the calibration measurements, and ordered by increasing axial position. All images are then co-registered using image cross-correlation. Next, the above mentioned spectral bleed-through is corrected by subtracting their intensities with corresponding spectral weighting factors from each color channel.

For reconstructing the 3D information of our volumetric image stacks, we applied 20 iterations of a 3D Lucy-Richardson algorithm [20, 21] (Matlab function *deconvlucy*) using pre-calculated point spread functions (PSFs). These 3D PSFs are calculated using the wave-optical theory of Wolf and Richards [22, 23]. Before applying the Richardson-Lucy deconvolution, the bottom and top image planes are mirrored and padded to each side of the image stack in order to reduce the boundary artifacts [24]. Finally, all three color stacks are cropped to a common axial range. The cropped volume is nearly 95% of the initially acquired volume which demonstrates the small axial chromatic aberration of our imaging system.

### 2.3. Cell culture, transfection, and immunostaining

COS-7 cell line CRL-1651™ was purchased from American Type Culture Collection. All cells were cultured in DMEM supplemented with 10% fetal calf serum (FCS), 2 mM L-glutamine, 1 mM sodium pyruvate, 100 U/ml penicillin / streptomycin in a humidified 5% CO_2_ atmosphere at 37 °C. Cells were tested negative for mycoplasma contamination.

For fixed cell samples, cells were grown on glass cover slips for 24 h, fixed in 3.6% formaldehyde in phosphate buffered saline (PBS) for 15 minutes, and washed with PBS. Then, cells were permeabilized with 0.5% Triton X-100 in PBS for 10 minutes and blocked with 3% BSA in PBS for 30 minutes at room temperature. Next, cells were incubated with rabbit monoclonal anti-vimentin antibody (SP20, Invitrogen, 1:500) in 3% BSA in PBS for one hour. After washing with PBS, cells were incubated with FITC-conjugated goat anti-rabbit IgG (ab6717, Abcam, 1:1000) together with 250 nM MitoTracker Deep Red FM (M22426, Invitrogen), and 100 nM phalloidin-Atto 550 (Atto-Tec GmbH) in 3% BSA in PBS for one hour. After staining, the cells were washed with PBS and rinsed with distilled water. The samples were mounted on glass slides using Fluoroshield (F6182, Sigma-Aldrich).

For live-cell measurements, cells were grown on glass coverslips for 24 h and then transfected with both mCherry-lamin B1-10 and GFP-connexin-43 using ViaFect (Promega). The mCherry-lamin B1-10 was a gift from Michael Davidson (Addgene plasmid #55069). GFP-tagged rat connexin-43 was generated earlier [25]. Staining of mitochondria and live-cell imaging was performed about 12 hours after transfection. For this purpose, 500 mM MitoTracker Deep Red FM was added directly to culture medium for 5 min at 37 °C. Then, coverslips were mounted into self-made microscope imaging chambers, and cells were supplied with fresh culture medium. During imaging, cells were kept in a humidified 5% CO_2_ atmosphere at 37 °C.

## 3. Results

We first validated the volumetric three-color imaging capability of our system by imaging of fixed COS-7 cells with three different fluorescently labelled intra-cellular structures: vimentin, actin, and mitochondria. Fig. 3 shows transverse slices as well as a lateral views of the imaged 40 × 40 × 2.5 μm volume, where vimentin is shown in cyan, actin in yellow, and mitochondria in red. An animated 3D representation of the imaged volume is shown by the supplemental movie ‘Video1’.

**Fig. 3.**
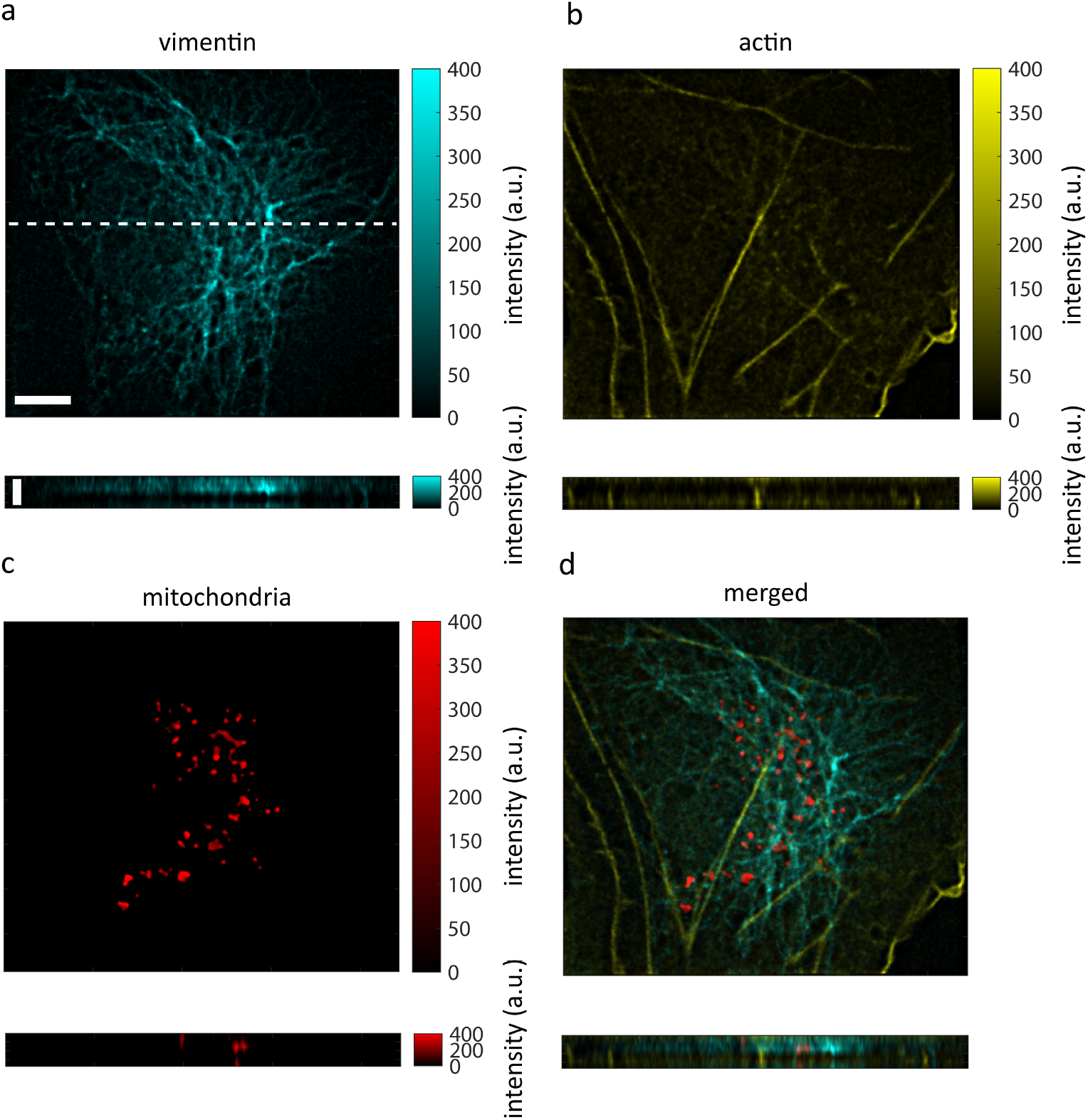
Instant 3D three-color imaging of a fixed COS-7 cell. (a) vimentin (stained with FITC-conjugated goat anti-rabbit IgG) imaged through the green channel in a *xy* section (upper image) and a *xz* section (lower image) along the white dashed line highlighted in the transverse image. (b), (c) the same views as (a) for actin (stained with phalloidin-Atto 550) and mitochondria (stained with MitoTracker deep Red) imaged respectively imaged through the orange and deep red channels. (d) Overlay of the three components. Scale bars in upper and lower panels in (a) indicate lengths of 5 μm and 2 μm, respectively.

Fig. 4 shows volumetric multi-color images of connexin (in cyan), nuclei (in yellow), and mitochondria (in red) in a living COS-7 cell at five different time points. Image acquisition time for each multi-color volume was 100 ms. One can observe both the lateral and axial motion of connexin and mitochondria over time (see also supplemental movies ‘Video2’, ‘Video3’, and ‘Video4’). Due to the highly dynamic behaviour of connexin, the median intensity over all frames is subtracted from each image to enhance the signal-to-noise ratio before deconvolution.

**Fig. 4.**
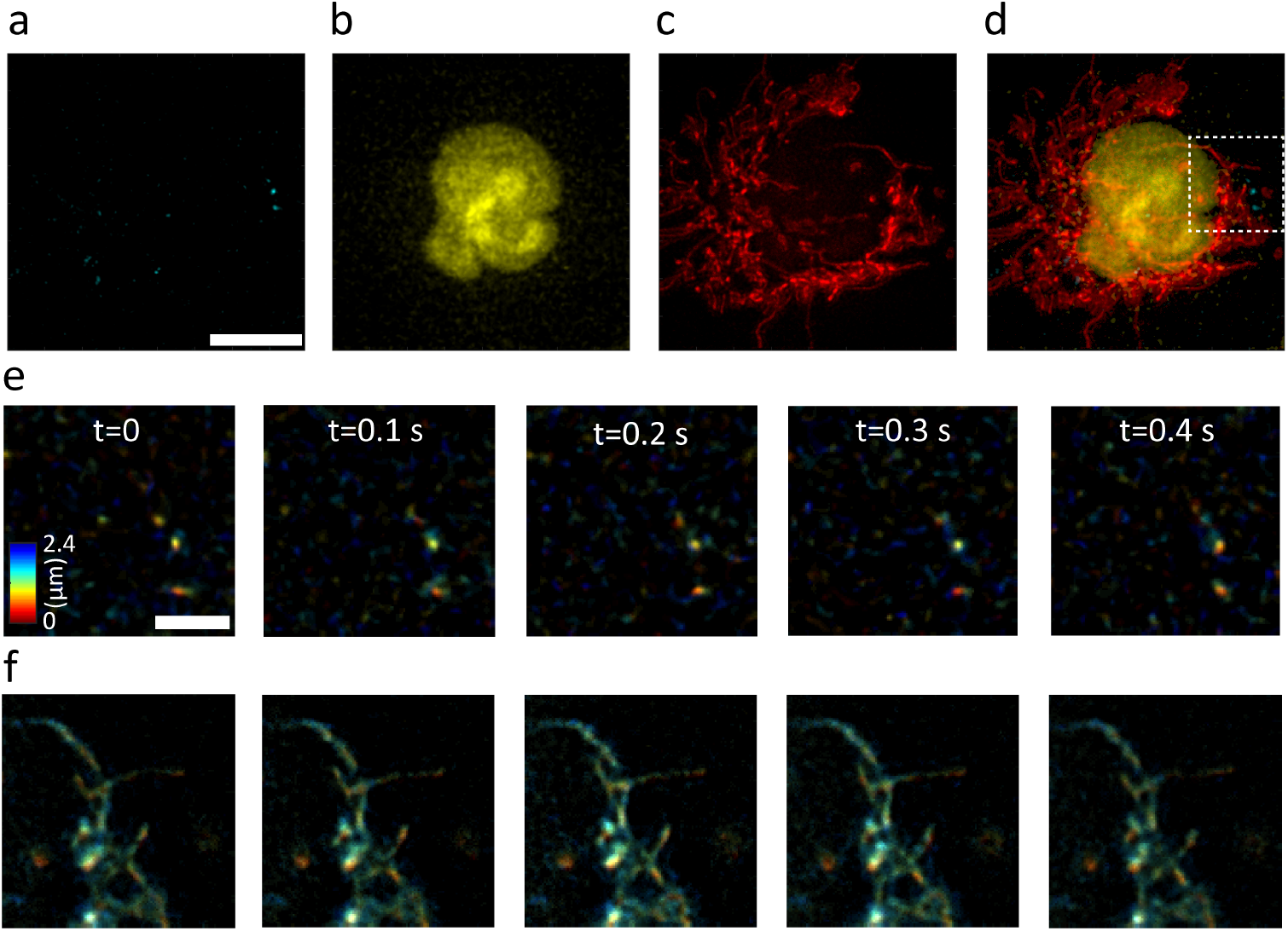
Instant 3D three-color live cell imaging of a COS-7 cell. Maximum intensity projection of (a) connexin (stained with GFP) imaged through the green channel, (b) nuclei (stained with mCherry) imaged through the orange channel, mitochondria (stained with MitoTracker deep red) imaged through the deep red channel, and (d) the merged image. (e) Time series of connexin dynamics indicated in the white dashed square in (d) in five frames where colors encodes the height of sample. (f) the same time points as (e) but showing the mitochondria dynamics. Scale bars in (a) 10 μm and in (e) 3 μm.

## 4. Discussion

We presented an easy to realize multi-plane three-color wide-field microscope that is capable of recording images with frame rate of several hundred three-color volume images per second (determined solely by the image acquisition speed of the used sCMOS cameras). We described in detail the setup, its calibration, and image processing routines. Our system exploits sCMOS cameras more efficiently than other existing multi-plane single color systems [12, 15, 19] because it employs nearly three forth of all pixels available in both sCMOS cameras. Our microscope exploits only commercially available components (most prominently optical dichroic and band pass filters, and a commercial multi-plane beam splitter). We demonstrated the performance of this microscope by imaging multi-color labelled fixed and live cells. It should be also mentioned that the axial range of volume imaging of our system is determined by the magnification. Thus, this range can be extended (or reduced) by appropriately changing the image magnification.

A potential future extension of our microscope could be its combination with super-resolution optical fluctuation imaging (SOFI), extending the work of ref. [26] to three-color imaging. Another important extension could be its combination with structured illumination microscopy (SIM), by using appropriate structured illumination in the excitation path. This would not only increases the lateral resolution of our system, but also significantly improves the optical sectioning which is, at the moment, only achieved indirectly via Richardson-Lucy deconvolution.

